# Global Transcriptomics of Congenital Hepatic Fibrosis in Autosomal Recessive Polycystic Kidney Disease using PCK rats

**DOI:** 10.1101/2023.01.19.524760

**Authors:** Satyajeet Khare, Lu Jiang, Diego Paine Cabrara, Udayan Apte, Michele T. Pritchard

**Author notes:** **Co-corresponding authors:** Michele T. Pritchard, PhD, Department of Pharmacology, Toxicology and Therapeutics, University of Kansas Medical Center, 3901 Rainbow Blvd, MS1018, Kansas City, KS 66160, Udayan Apte, PhD, DABT, Department of Pharmacology, Toxicology and Therapeutics, University of Kansas Medical Center, 3901 Rainbow Blvd, MS1018, Kansas City, KS 66160.

## Abstract

Congenital hepatic fibrosis / Autosomal recessive polycystic kidney disease (CHF/ARPKD) is an inherited neonatal disease induced by mutations in the *PKHD1* gene and characterized by cysts, and robust pericystic fibrosis in liver and kidney. The PCK rat is an excellent animal model which carries a *Pkhd1* mutation and exhibits similar pathophysiology. We performed RNA-Seq analysis on liver samples from PCK rats over a time course of postnatal day (PND) 15, 20, 30, and 90 using age-matched Sprague-Dawley (SD) rats as controls to characterize molecular mechanisms of CHF/ARPKD pathogenesis. A comprehensive differential gene expression (DEG) analysis identified 1298 DEGs between PCK and SD rats. The genes overexpressed in the PCK rats at PND 30 and 90 were involved cell migration (e.g. *Lamc2, Tgfb2*, and *Plet1*), cell adhesion (e.g. *Spp1, Adgrg1*, and *Cd44*), and wound healing (e.g. *Plat, Celsr1, Tpm1*). Connective tissue growth factor (*Ctgf*) and platelet-derived growth factor (*Pdgfb*), two genes associated with fibrosis, were upregulated in PCK rats at all time-points. Genes associated with MHC class I molecules (e.g. *RT1-A2*) or involved in ribosome assembly (e.g. *Pes1*) were significantly downregulated in PCK rats. Upstream regulator analysis showed activation of proteins involved tissue growth (MTPN) and inflammation (STAT family members) and chromatin remodeling (BRG1), and inhibition of proteins involved in hepatic differentiation (HNF4α) and reduction of fibrosis (SMAD7). The increase in mRNAs of four top upregulated genes including *Reg3b, Aoc1, Tm4sf20*, and *Cdx2* was confirmed at the protein level using immunohistochemistry. In conclusion, these studies indicate that a combination of increased inflammation, cell migration and wound healing, and inhibition of hepatic function, decreased antifibrotic gene expression are the major underlying pathogenic mechanisms in CHF/ARPKD.

## Introduction

Autosomal recessive polycystic kidney disease (ARPKD) is a hereditary disorder caused by mutations in *PKHD1* ^1^. It is a ciliopathy where the growth and function of the primary cilium is affected on several types of epithelial cells, including renal tubule cells and biliary tree cholangiocytes, leading to cystogenesis. *PKHD1* codes for the protein fibrocystin, which is expressed on the primary cilium in the epithelial cells and mutations in *PKHD1* disrupt primary cilium function^2, 3^. The clinical spectrum is widely variable with multiple organs affected but primarily the kidneys and the liver. Kidney disease is characterized by tubule dilation and focal cyst development in collecting ducts ^4, 5^. Liver disease in ARPKD is a characterized by congenital hepatic fibrosis (CHF), which involves development of segmental dilations of bile ducts accompanied by periportal fibrosis ^6^. CHF causes significant morbidity and mortality in ARPKD patients who develop portal hypertension ^7^. Although the mechanisms driving hepatic cyst expansion are not well characterized in human CHF/ARPKD, different factors, such as cholangiocyte hyperproliferation and fluid secretion, and perhaps altered cell-cell and cell matrix biomechanics ^8^ may contribute to the progression of cysts ^9^. One of the best-characterized animal models of CHF/ARPKD is the PCK rat, derived from a colony of Sprague-Dawley (SD) rats at Charles River, Inc ^10^. The PCK rat carries a gene mutation that is orthologous to human *PKHD1* ^11^. As a spontaneously occurring, inherited model of ARPKD, PCK rat hepatorenal fibrocystic disease bears a striking resemblance to human ARPKD ^12^.

The mechanisms, especially those underlying gene expression involved in CHF/ARPKD pathogenesis, are not completely known. We previously proposed a ‘pathogenic triumvirate’ to describe the three pathogenic components in CHF/ARPKD, which includes cyst growth, inflammation, and fibrosis ^13^. Using RNA-seq analysis to analyze liver samples from SD rats and PCK rats at different time points after birth, we found that differentially expressed genes, transcription factors, and canonical pathways in PCK can be categorized into one of these three components. In addition, we identified several genes and transcription factors related to liver development and metabolism in PCK rats that may contribute to the pathogenesis of CHF/ARPKD and confirmed a subset of the top upregulated genes in PCK rat tissue. Overall, our study provides a comprehensive characterization of gene expression profile in CHF/ARPKD. By identifying potential regulators of the ‘pathogenic triumvirate’ and other possible contributors of CHF/ARPKD, we propose several molecular targets of the disease that can be explored in further mechanistic studies and with the overall aim of developing novel therapeutic strategies.

## Materials and methods

### Animals

Male and female PCK rats and Sprague-Dawley (SD) rats (control strain for PCK) were purchased from Charles River and housed in an AAALAC-accredited vivarium at the University of Kansas Medical Center (KUMC) and used to generate rat pups for these studies. All animal studies were approved by the KUMC Institutional Animal Care and Use Committee (IACUC) and were performed in accordance with The Guide for the Care and Use of Laboratory Animals (8^th^ Edition).

### Liver tissue collection and storage

Rat pups were aged to postnatal day (PND)15, 20, 30 or 90 and then euthanized by isoflurane inhalation overdose. One euthanized, the livers were excised, dissected into separate lobes, and further cut into smaller pieces. Some liver pieces were snap frozen in liquid nitrogen and stored at -80°C for RNA sequencing analysis while other liver pieces were placed into histological cassettes and submerged in 10% neutral buffered formalin for histological analysis. Paraffin embedded liver sections were used for H&E and Mason’s Trichrome staining.

### Sample preparation and RNA isolation

Liver samples were collected from SD and PCK rats at postnatal day (PND) 15, 20, 30, and 90 (n = 3 each time point for each genotype). Both male and female rats were included. Previously snap-frozen liver pieces (20 – 30 mg) were transferred to 1.5 mL of RNAlater-ICE (Ambion Inc., Austin, TX) and stored at −80 °C overnight before RNA isolation. Total RNA was isolated with the RNeasy Mini Kit (Qiagen, Valencia, CA) after homogenization using a bead homogenizer with lysing matrix D homogenization tubes (MP Biomedicals Fast Prep 24, Solon, OH). The total RNA concentration and purity were determined using a NanoDrop microvolume spectrophotometer (Thermo Fisher Scientific, Lenexa, KS) apparatus.

### RNA library preparation and data collection

Total RNA (500 ng) was used to construct the Stranded mRNA-Seq library. Briefly, the total RNA fraction was fragmented, reverse transcribed into cDNA, and ligated with the appropriate indexed adaptors using the TruSeq Stranded mRNA LT Sample Preparation Kit (Illumina, San Diego, CA). The quality of library was validated by an Agilent Bioanalyzer. Library quantification was measured using the Roche Lightcycler96 with FastStart Essential DNA Green Master Mix. Library concentrations were adjusted to 4 nM and pooled for multiplexed sequencing. Libraries were denatured and diluted to the appropriate pM concentration followed by clonal clustering onto the sequencing flow cell using the TruSeq Paired-End (PE) Cluster Kit v3-cBot-HS. The clonal clustering procedure is automated using the Illumina cBOT Cluster Station. The clustered flow cell was sequenced on the Illumina HiSeq 2500 Sequencing System using the TruSeq SBS Kit v3-HS. Sequence data is converted to Fastq format from Bcl files and de-multiplexed into individual sequences for further downstream analysis.

### Bioinformatic analysis for RNA-Seq

Quality of sequence reads was confirmed using FASTQC. Reads were aligned to the reference genome Rnor 6.0 from Ensembl^14^ using HiSAT2^15^. The bam files generated were used to create an expression matrix using featureCounts^16^. Differential expression analysis was performed on the count matrix using DESeq2^17^. Differentially expressed genes identified were subjected to enrichment analysis using clusterProfiler^18^. Expression plots were generated using normalized CPM values in R^19^.

Pathway analysis was performed using Ingenuity Pathway Analysis (IPA, Ingenuity Systems, www.ingenuity.com). Differential expression was determined by absolute fold change > 1.5 and a q-value (the p-value adjusted for multiple hypothesis correction using the Benjamini-Hochberg procedure) less than or equal to 0.05. IPA upstream analysis was performed to predict the activated and inhibited upstream regulators using activation of z-score algorithm. The activation Z-score is applied to quantify the significance of the concordance between the direction of change of genes (up-/down-regulated) in a gene dataset and the established direction of change (derived from literature) of genes associated with a transcriptional regulator. A z-score ≥ 2 or ≤ -2 is considered significantly activated or inhibited, respectively. P value of overlap reflects that the overlap between the experimental dataset and a given transcriptional regulator is due to a random chance. An overlap p-value ≤0.05 was considered significant.

### Statistical Analysis of non-RNAseq data

All non-RNA-seq data were expressed as mean ± standard error of the mean. Student’s t-test or ANOVA with Tukey’s post-hoc were used for analyses with a p-value <0.05.

### Data Availability

The datasets generated for this study can be found in NCBI SRA, PRJNA799864

## Results

### Progressive pericystic fibrosis in PCK rats

PCK rats are a variant of the Sprague Dawley (SD) rat strain which harbors mutations in the *Pkhd1* gene, the gene orthologous to that which causes ARPKD in humans. Previous studies extensively characterized this model and demonstrated that PCK rat mimic the pathophysiological characteristics of human CHF/ARPKD. These rats develop extensive hepatic cysts, which are present at birth and progressively increase in number and size with age^11^. This results in extensive hepatomegaly in PCK rats as compared to SD rats demonstrated by significantly higher liver weight to body weight ratio, which is evident by PND10 and continues to increase till PND90 (Fig. 1A) Additionally, extensive pericystic fibrosis is evident in the PCK rats as demonstrated using Masson’s trichrome staining (Fig. 1B). The kidney and liver disease in the PCK rats becomes progressively worse necessitating euthanasia around 6 months of age. These characteristics make PCK rats an ideal model to investigate mechanisms of hepatic cystogenesis and fibrosis in CHF/ARPKD.

**Figure 1.**
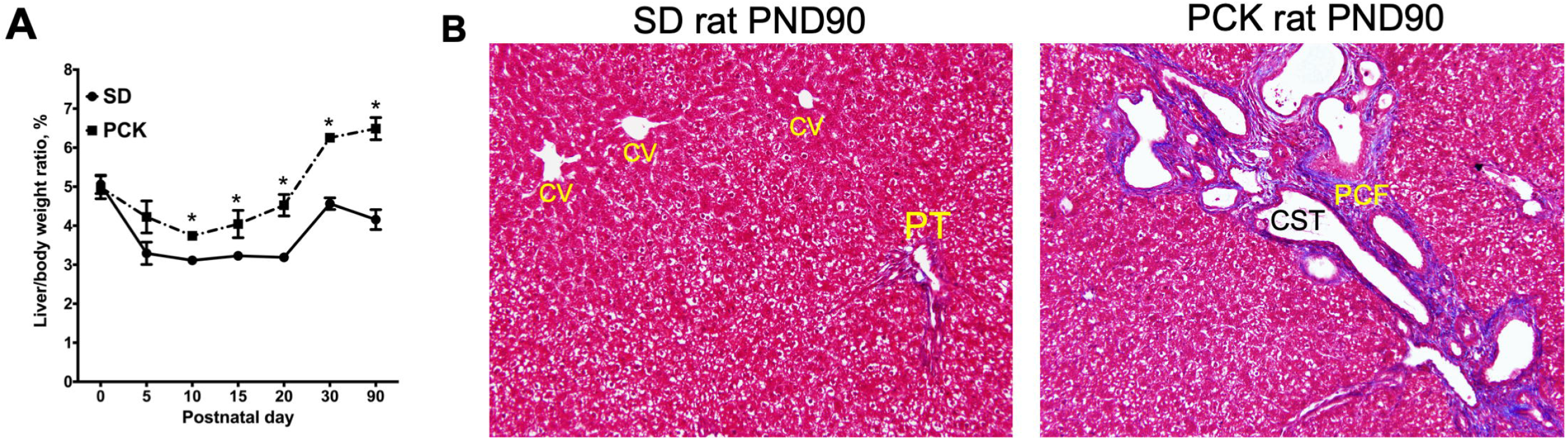
Histopathological features of ARPKD/CHF. Representative photomicrographs of Masson’s trichrome stained liver sections from (A) SD rats and (B) PCK rat at PND90. Blue staining localizes collagen. CV, central vein; CST, hepatic cyst; PCF, pericystic fibrosis; PT, portal triad area.

### RNAseq analysis correlated with progressive worsening of CHF/ARPKD

We performed global whole liver gene expression analysis using three individual SD and PCK rats at 3 time points (PND15, 20, 30 and 90, total n=24]. Differentially expressed gene (DEG) analysis showed a progressive increase in number of genes significantly different between SD and PCK rats with highest difference in PND90 (Table 1). Principle component analysis (PCA), distance matrix analysis and correlation plots revealed that the biological replicates at each time point clustered together indicating excellent quality of the samples (Fig. 2A and Supplementary Fig. 1). PCA further showed a time dependent change in hepatic gene expression between control SD livers and PCK livers. SD-PND15 and SD-PND20 samples clustered together with PCK-PND15 and PCK-PND20 samples suggesting relatively less overall difference in hepatic gene expression during these early time points. A significant shift in clustering was observed at PND 30 which continued further at PND 90. PCA showed that SD-PND30 samples clustered separately from SD-PND90, an expected result due to hepatic maturation between 1 and 3 months of age. Interestingly, the PCK-PND30 and PCK-PND90 samples clustered separately from each other and from SD rat samples at the same time points indicating both age related changes and changes due to disease progression.

**Table 1.** Differential gene expression* in PCK rats as compared to SD rats

| **PND** | **Upregulated** | **Downregulated** |
| --- | --- | --- |
| 15 | 620 | 667 |
| 20 | 1860 | 2174 |
| 30 | 2732 | 2767 |
| 90 | 3208 | 3279 |

\* Based on adj. p-value cut-off of 0.01. No fold change cut-off was applied.

**Figure 2.**
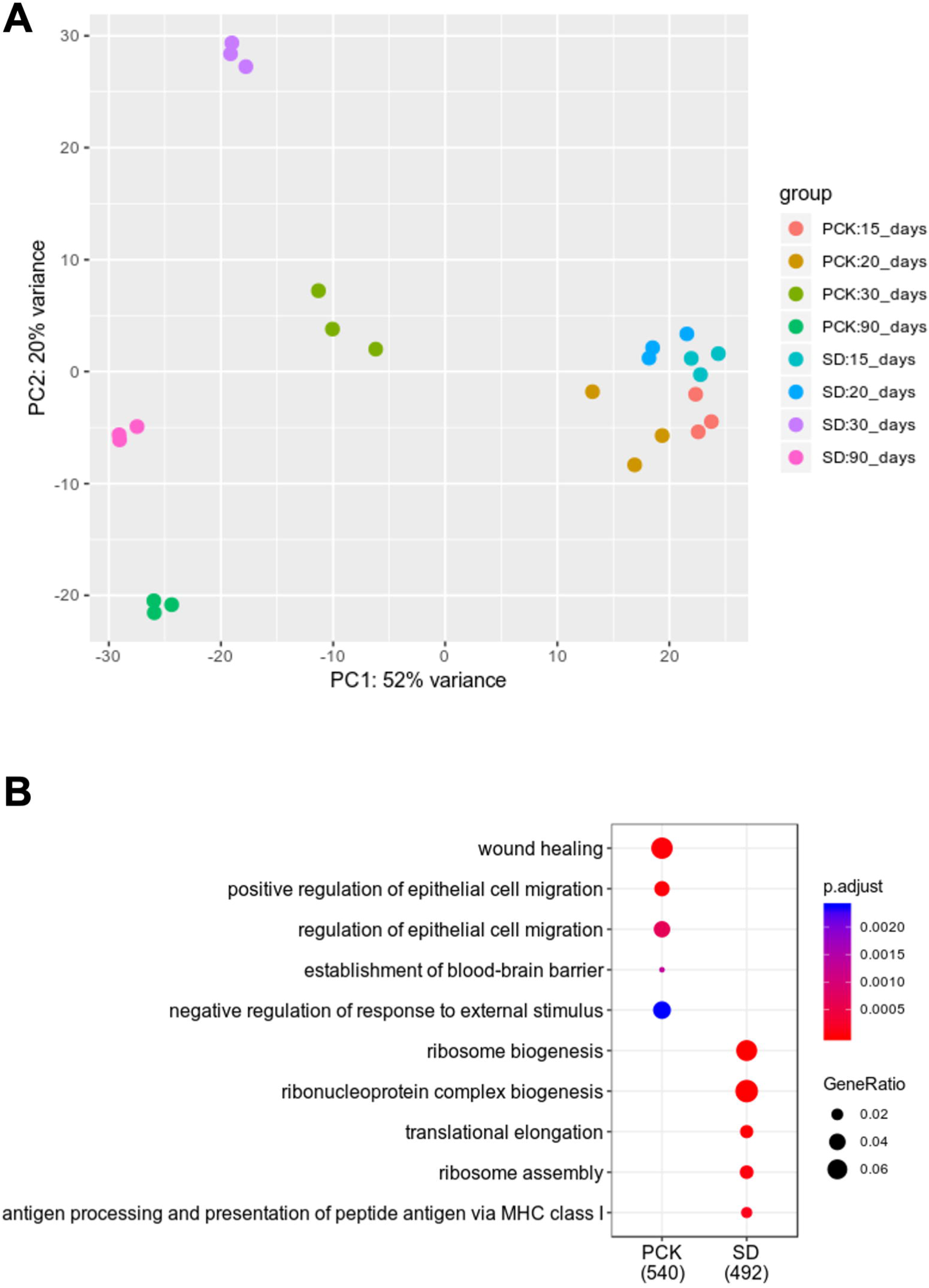
Characteristics of whole liver RNAseq data. (A) Principle component analysis revealed clustering of biological replicates withtin a group, suggesting good reproducibility. The samples are separated by age on the first principal component and genotype on the second principal component. This indicates more age related changes relative to gentoypte related changes in gene expression. Also, there are fewer differentially expressed genes in younger age liver samples (e.g. PND15) than older age samples (e.g. PND90). (B) Gene ontology analysis showing overall differential genes expression in SD and PCK rats

### Gene Ontology (GO) analysis reveals major categories of DEGs

We performed a differential gene expression analysis with adjusted P<0.01, which revealed that 1298 genes were differentially expressed between SD and PCK rats when all time points were considered together. We used this strategy to capture major changes due to both disease progression (time) and genotype. Gene ontology analysis revealed a significant activation of genes involved in wound healing, cell migration, and cell adhesion while a significant downregulation of genes involved in ribosomal biogenesis, and antigen presentation in PCK rats as compared to SD rats (Fig. 2B).

The genes associated with wound healing could be further clustered in two subclusters which changed expression pattern over the time course (Fig. 3A). The first cluster contained (outlined by a red box) genes such as *Plat, Wnt7a, Tpm1* and *Pdgfb* and showed higher expression in PCK rats at PND 15 and PND 20 with a moderate reduction at PND 30 and PND 90 but were expressed still higher than SD rats at the respective time points. The second cluster (outlined by a blue box) genes such as *Crp, P2ry2, Tgfbr2*, and *Mmp12* showed an overall lower expression as compared to the first cluster in PCK rats. However, expression of these genes was moderately higher in PCK rats when compared to SD rats at all time points except PND 30.

**Figure 3.**
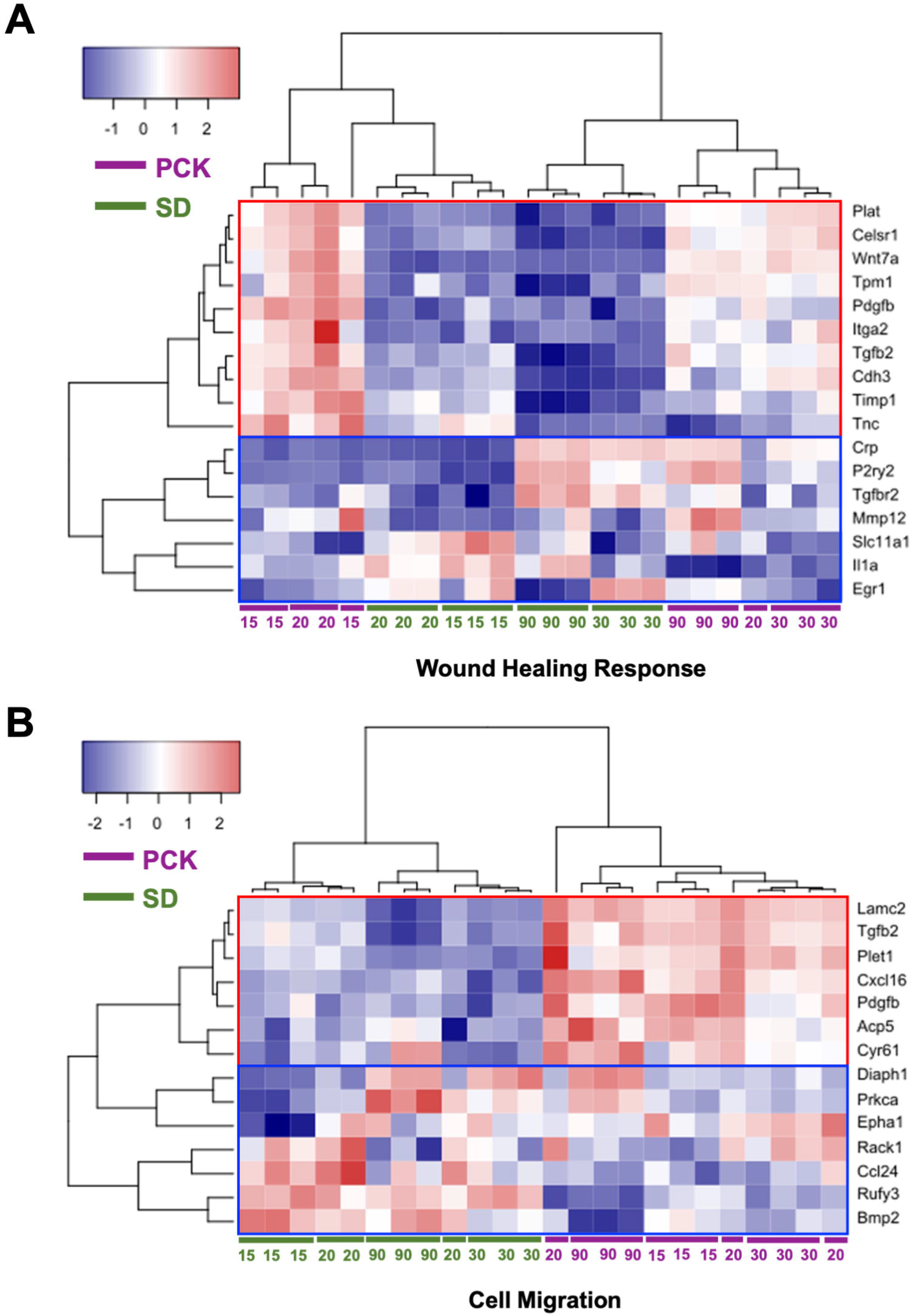
Genes involved in wound healing and cell migration are induced in PCK rats. (A) Heatmap showing significant changes in genes involved in wound healing response in PCK rats over time. Many of these genes were associated with change in extracellular matrix including *TGF-beta, Igta1, Pdgfb, Cdh3, Timp1* and *Mmp12*. (B) Heatmap of differntially expressed genes associated with cell migration including *CxCl16, Tgfb2, Lamc2* and *Cyr61* in PCK rats. Postnatal age (PND15, 20, 30, 90) is indicated at the bottom of the heatmap where SD rat data are labeled in green text and PCK data are labeled in purple text. Red squares indicate upregulated genes, blue squares indicate down regulated genes; color intensity indicates magnitude of change. The red and blue outlines in each heatmap indicate gene expression subclusters based on relative expression between PCK and SD rats over time. See text for details.

The genes Involved in cell migration also segregated in two subclusters and illustrated complete segregation of genes between genotypes suggesting disease specific regulation (Fig. 3B). The first subcluster (outlined by a red box) containing genes such as *Lamc2, Tgfb2, Cxcl16, Acp5* were significantly higher in PCK rats at all time points whereas the second gene cluster (outlined by a blue box), which included *Diaph1, Prkca, Rufy3* and *Bmp2*, showed an overall lower expression in PCK rats at most of the time points as compared to SD rats.

A significant number of DEGs were categorized as involved in cell adhesion by GO analysis, which formed 3 approximate subclusters (Fig. 4). The first cluster (outlined by a red box) containing genes such as *Spp1, Cd44, Pkp1*, and *Mybpc3*, showed significantly higher expression in PCK rats at all time points as compared to SD rats. A second cluster (outlined by a blue box) containing genes such as *F11r, Cd47, Itgae* and *Cadm1*, showed a temporal change between the groups. Most of the genes in this group showed significantly higher expression in PCK rats at PND 15 and 20 as compared to SD rats. A moderate increase in expression of these genes was observed at PND 30 and 90 in SD rats but it was still significantly lower than the expression in PCK rats at PND30 and especially at PND 90. A third cluster (outlined by a black box) containing genes such as *Lrfn3, Cd99l2, Pnn*, and *Cd4* showed temporal variation during between PND 15-20 and PND 30-90 time points. In SD rats, these genes were higher at PND 15 and 20 but showed decreased expression at PND 30 and 90. In PCK rats, expression of these genes was higher at PND 15 and 20 as compared to PND 30 and 90 but was significantly lower than SD rats at the same time points.

**Figure 4.**
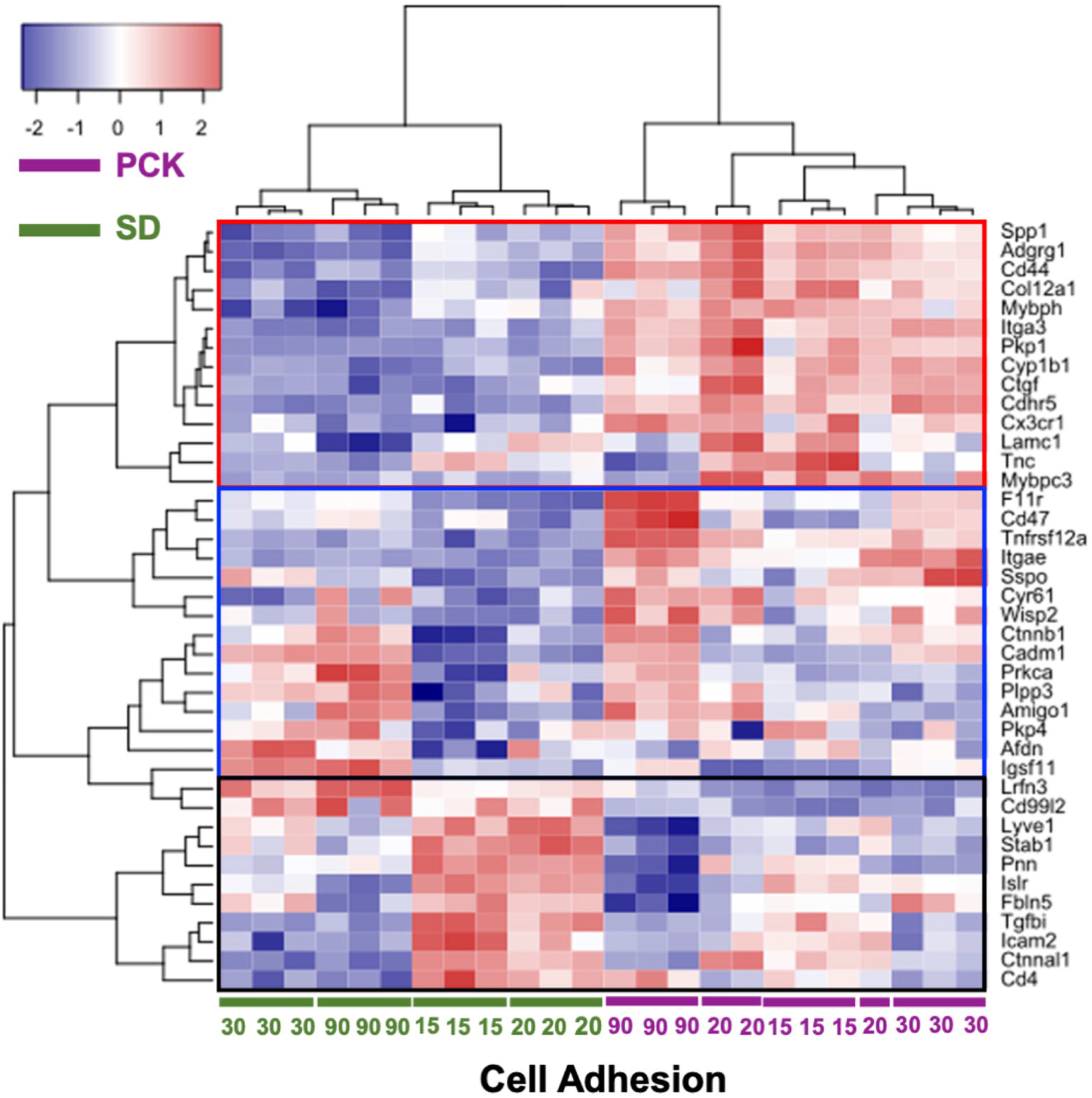
Dysregulated cell adhesion gene expression expression in PCK rats. Heatmap showing significant induction of several cell adhesion genes including *CD44, Spp1, Itga3, Col12a1* and *Icam* in PCK rats (subcluster with red outline). Subclusters outlined in blue and black contain genes with temporal changes over time or that are mostly down regulated in PCK rats, respectively. Postnatal age (PND15, 20, 30, 90) is indicated at the bottom of the heatmap where SD rat data are labeled in green text and PCK rat data are labeled in purple text. Red squares indicate upregulated genes, blue squares indicate down regulated genes; color intensity indicates magnitude of change. The red, blue and black outlines in the heatmap indicate gene expression subclusters based on relative expression between PCK and SD rats over time. See text for details.

Two categories of genes showed significant downregulation in PCK rats relative to SD rats including those involved in antigen presentation (Fig. 5A; *RT1-A2, RT1-CE10* etc.) and those involved in ribosomal biogenesis (Fig. 5B; *Pes1, Rsp28* etc.). The genes involved in antigen presentation showed a significantly lower in expression at PND 15 and 20 in PCK rats than the SD rats, showed a moderate increase in expression at PND 30 and 90 as compared to PCK-PND15/20 but were still lower in expression than the SD-PND 30/90 groups (Fig. 5A). The genes involved in ribosomal biogenesis showed an extensive variation in their expression patterns between the time points and genotypes but were overall lower in PCK rats as compared to SD rats (Fig. 5B).

**Figure 5.**
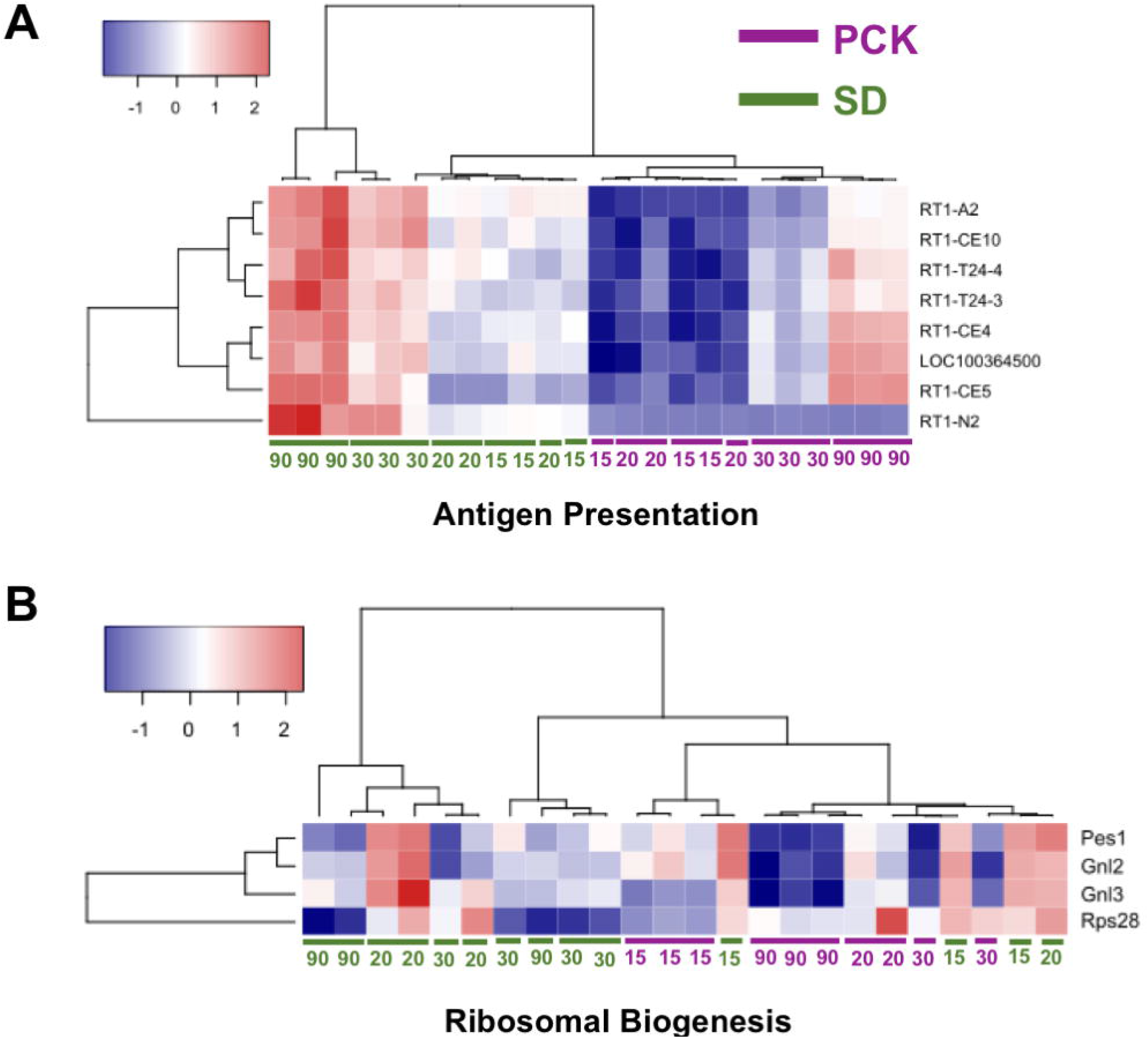
Downreglated genes in PCK rats. Heatmap showing significant decrease in expression of genes associated with (A) antigen presentation and (b) ribosomal biogenesis in PCK rats. Postnatal age (PND15, 20, 30, 90) is indicated at the bottom of the heatmap where SD rat data are labeled in green text and PCK data are labeled in purple text. Red squares indicate upregulated genes, blue squares indicate down regulated genes; color intensity indicates magnitude of change.

### Identification of upstream regulators and canonical pathways in PCK rats

To further investigate the global gene expression data and identify key players, we performed an upstream regulator analysis using Ingenuity Pathway Analysis (IPA) software. The upstream regulators were predicted based on the IPA database. The activated upstream regulators (Table 2) showed some consistently upregulated molecules such as MTPN (myotrophin), which was predicted to be activated at all four time points in PCK rats. SMARCA4, which codes for the protein BRG1 was activated at PND 20 and 30. Interestingly, various members of STAT family of transcription factors were activated at all four time points. (PND15- STAT5B, PND20- STAT4, PND30- STAT3 and STAT1, and PND90- STAT3 and STAT4).

**Table 2.** Changes in Upstream regulators in PCK rats

| Activated Upstream Regulators |  |  |  |  |  |  |  |
| --- | --- | --- | --- | --- | --- | --- | --- |
| 15 |  | 20 |  | 30 |  | 90 |  |
| Name | Z-score | Name | Z-score | Name | Z-score | Name | Z-score |
| CTNNB1 | 2.782 | RB1 | 3.359 | MTPN | 3.44 | STAT3 | 4.599 |
| MTPN | 2.433 | STAT4 | 3.174 | IRF7 | 3.01 | FOXO1 | 4.186 |
| TCF7L2 | 2.167 | SMARCA4 | 3.022 | SMARCA4 | 2.847 | STAT4 | 4.016 |
| STAT5B | 2.000 | MTPN | 2.728 | STAT3 | 2.641 | MTPN | 3.501 |
|  |  | ARNT | 2.513 | STAT1 | 2.629 |  |  |
| Inhibited Upstream Regulators |  |  |  |  |  |  |  |
| 15 |  | 20 |  | 30 |  | 90 |  |
| Name | Z-score | Name | Z-score | Name | Z-score | Name | Z-score |
| CBX5 | -2.236 | KDMSA | -2.530 | HNF4A | -3.029 | HNF4A | -2.821 |
|  |  | CBX5 | -2.309 | ATF4 | -2.94 | SMAD7 | -2.668 |
|  |  |  |  | CBX5 | -2.84 | ZFP36 | -2.559 |
|  |  |  |  | NFE2L2 | -2.775 | SNAI1 | -2.509 |
|  |  |  |  | SMAD7 | -2.691 | HDAC5 | -2.236 |

Th upstream regulators predicted to be inhibited in PCK rats (Table 2) included CBX5, also called HP1α, is a gene silencer which primarily works via H3K9 methylation and was predicted to be inhibited at PND15, 20, and 30. HNF4α, the master regulator of hepatocyte differentiation, was also predicted to be downregulated at PND30 and PND90. Finally, inhibition of SMAD7, an inhibitory SMAD which attenuates TGFβ and activin mediated signaling, was inhibited at PND 30 and 90 in PCK rats.

### Immunohistochemical confirmation of top upregulated genes

We performed a differential gene expression analysis to identify genes that were up or down regulated at all time points in PCK rats as compared to SD rats. The top five up and down regulated genes are listed in Table 3. Next, we performed immunohistochemical analysis to detect protein expression of the top four upregulated genes including *Reg3b, AOC1, TM2SF40*, and *CDX2* at PND30 (Fig. 6) and PND90 (Fig. 7) using paraffin embedded liver sections from SD and PCK rats. AOC1, CDX2 and Reg3b showed higher expression both in cyst wall epithelium and hepatocytes in PCK rats at both PND30 and PND90. TM2SF40 showed mainly nuclear staining pattern and was expressed at equal levels in SD and PCK hepatocytes. However, PCK rats also showed significant TM2SF40 expression in cyst wall epithelium. Further, at both PND30 and PND90, PCK rat hepatocytes also showed cytoplasmic expression of TM2SF40 in addition to nuclear expression, which was absent in SD rats. Interestingly, there very low to no staining of any of these proteins in the extracellular matrix-rich pericystic region, suggesting a preferential localization of proteins to the hepatic epithelial cell compartment.

**Table 3.** Top five up and down regulated genes in PCK rats at all time points

| **Gene Name** | **Fold Change** | **Gene Name** | **Fold Change** |
| --- | --- | --- | --- |
| Reg3b | 175.49 | Abhd3 | -101.76 |
| AOC1 | 72.31 | Pla2g4c | -72.09 |
| Tm4sf20 | 25.48 | Ly6al | -22.77 |
| Cdx2 | 10.30 | Mpped1 | -19.74 |
| Krt17 | 10.30 | Stfa2l1 | -5.21 |

**Figure 6.**
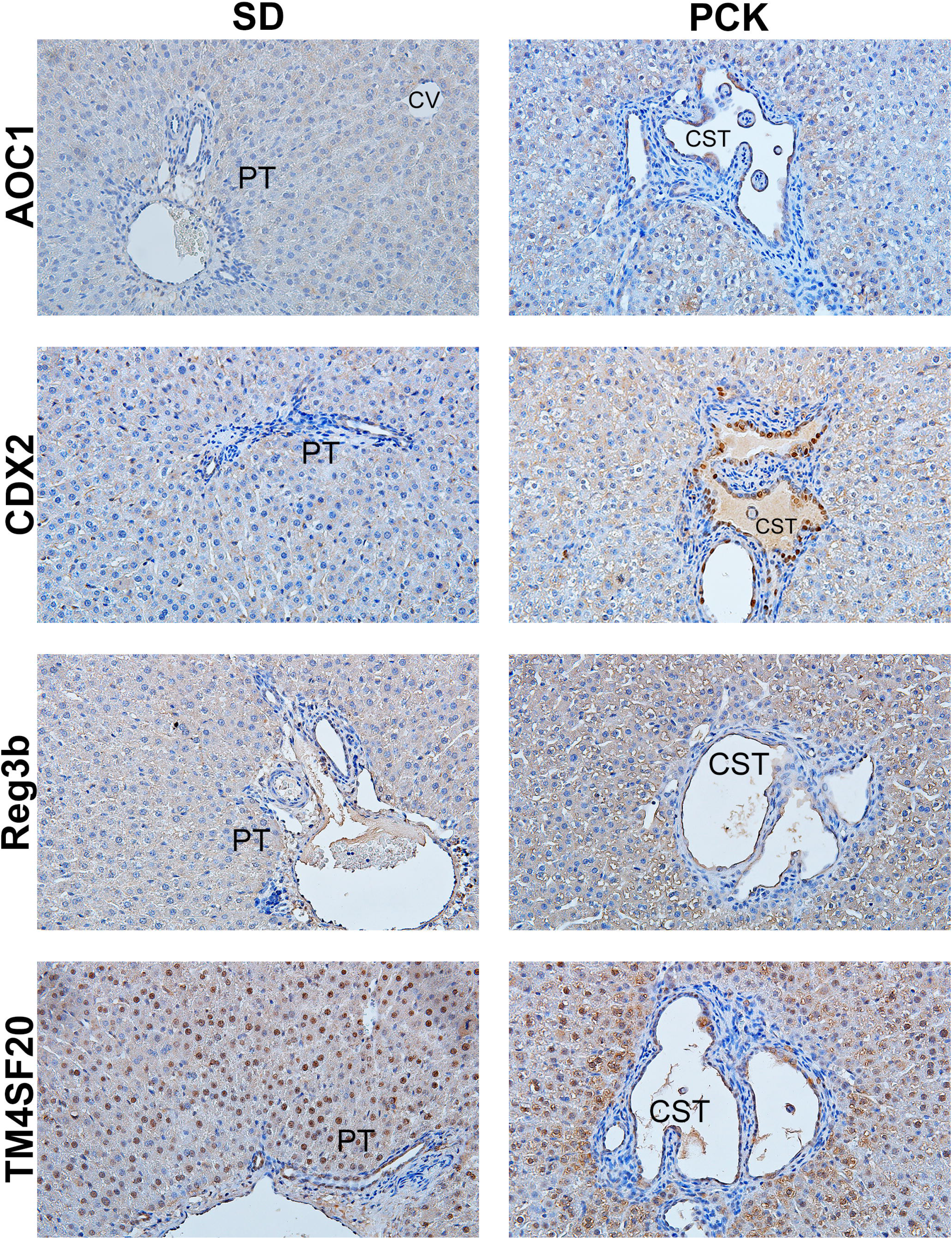
Immunohistochemical confirmation of top upregulated genes at PND 30. Representative photomicrographs (400x) of IHC for AOC1, CDX2, Reg3b and TM4SF20 was performed on formalin-fixed paraffin embedded liver sections from PND30 SD and PCK rats (n = ??) PT, portal triad; CV, central vein; CST, hepatic cyst.

**Figure 7.**
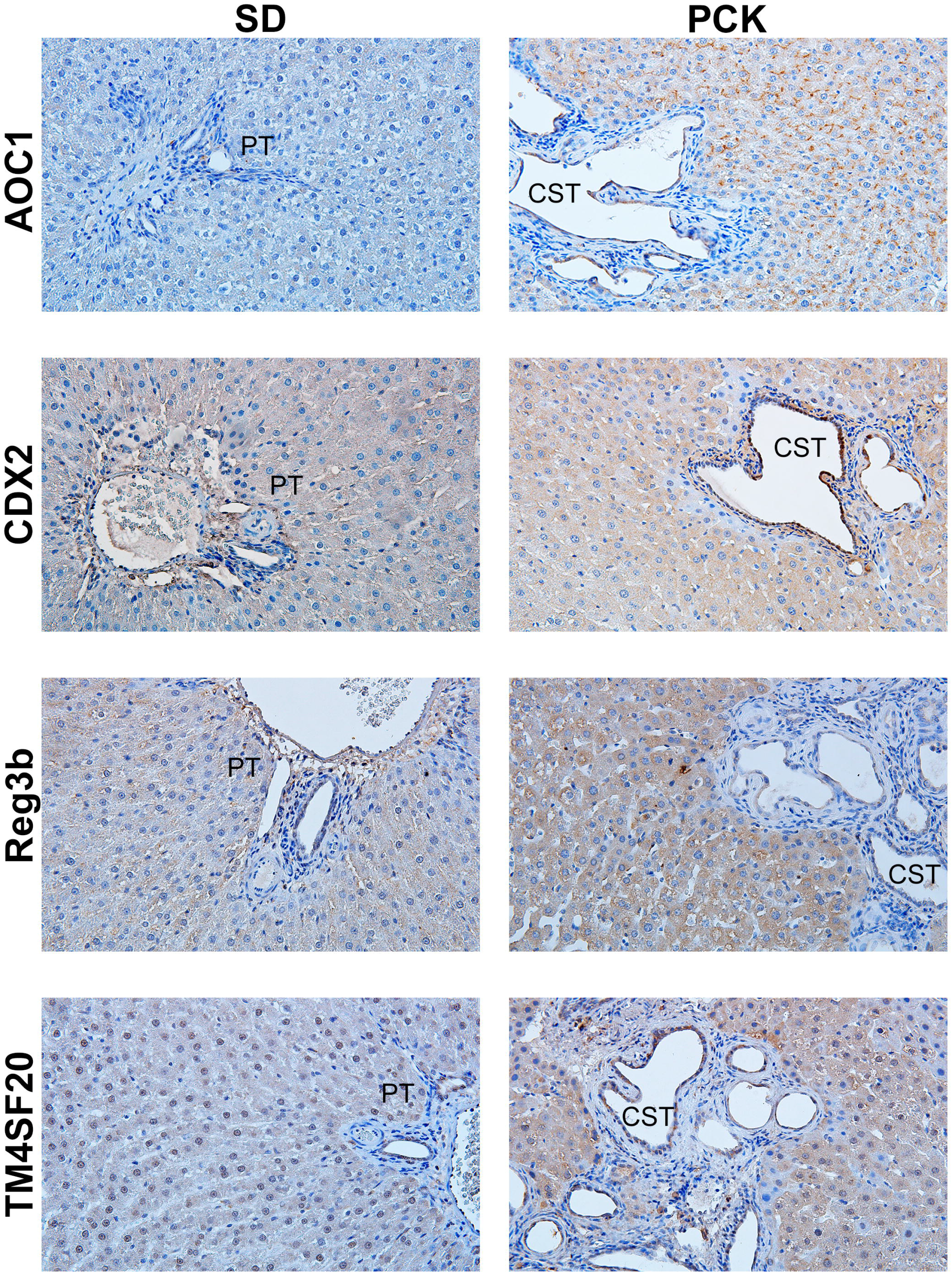
Immunohistochemical confirmation of top upregulated genes at PND90. Representative photomicrographs (400x) of IHC for AOC1, CDX2, Reg3b and TM4SF20 was performed on formalin-fixed paraffin embedded liver sections from PND90 SD and PCK rats. PT, portal triad; CV, central vein; CST, hepatic cyst.

## Discussion

CHF/ARPKD is a progressive genetic disease affecting mainly the kidneys and the liver. Although the phenotypes of CHF/ARPKD are well-characterized, the mechanisms driving these phenotypes in CHF/ARPKD are still incompletely understood. Previous studies indicate that a pathogenic triumvirate of inflammation, proliferation and fibrosis is at the center of progressive cyst development^20^. To reveal possible mechanisms, we performed RNA-Seq analysis in SD and PCK rats at various time points to compare the gene expression alterations associated with mutation in gene *PKHD1*. Our analysis focused on determining differential gene expression in developing (PND 15, 20, 30) and mature (PND 90) rats in both genotypes. This strategy is effective because it captures gene changes that occur as a function of hepatic maturation with time and those that occur due to presence of disease. A total of 1298 genes were significantly different between PCK livers and SD control livers with approximately 50% downregulated and other 50% upregulated. The principal component analysis indicated an age-related change in gene expression. This is interesting because it suggests that the normal hepatic maturation and differentiation process is disrupted in CHF/ARPKD. Rodent livers undergo a massive transcriptomic transformation during the first 6 weeks (42 days) of life during which the liver represses the embryonic gene expression program and induces the adult gene expression program. This differentiation process is combined with bursts of cell proliferation that aid in increasing the liver size to achieve the adult liver to body weight ratio. By 6 weeks of age, rodent livers look like adult livers both in terms of size and transcriptome (ref). Our analysis indicates that this normal process of size increase and hepatic maturation is severely dysregulated in PCK rats, and a growing number of transcriptomic changes occur leading to severe and rapid disease progression. PCA showed that up to PND20 there was a relatively small difference in gene expression between SD and PCK rats. By PND30 the two genotypes significantly separated indicating the ‘window of pathogenesis’ lies between PND15 and PND30. Extrapolating these findings to human condition will be both challenging and rewarding as the process of hepatic maturation is significantly more delayed requiring up to 2 years in humans^21^.

The gene ontology analysis of these data revealed a clear picture of disease progression involving disruption in cell migration and cell adhesion processes that ultimately produce a prolonged wound healing response. This has been observed in other fibrotic disorders in the liver and other tissues^22-24^. Loss of PKHD1 in PCK rat livers triggers an injury response that is not typical in the sense of structural damage or functional damage but seems to be associated with disruption of normal bile duct development. It is known that bile duct development continues during the early postnatal period where the biliary tree is expanding along with increasing liver size. It is plausible that loss of PKHD1 function in cholangiocytes along with other mechanisms induces cystogenesis, which is treated by the liver as a massive structural damage, perhaps due to micromechanical changes induced by expanding cysts, promoting a wound healing response and fibrosis. The gene expression profiles support this hypothesis as it shows increased expression of genes involved in cell migration and cell adhesion, both possible components of cyst formation and budding off from the biliary tree. The wound healing response, which seems to drive the fibrosis, is likely a response to increased micromechanical stiffness induced by these growing number of cysts. Whereas the exact signaling mechanisms remain to be investigated, our data provide several novel leads that were previously unknown.

The data on upstream regulators provides further evidence that the pathogenic triumvirate of proliferation, inflammation and fibrosis is at the center of the CHF/ARPKD disease processes^20^. One of the most highly and consistently predicted to be upregulated transcription factors as members of the signal transducers and activators of transcription (STAT) family including STAT1, STAT3, STAT4, and STAT5B, which are involved in growth and inflammatory processes. Previous studies have shown strong activation of STAT3, regulated by polycystin-1 (PC-1), the gene product of *PKD1* (mutated in ADPKD), in cyst-lining epithelial cells in mouse and human ADPKD ^25^.

Further, treatment with STAT3 inhibitors decreased cell proliferation in human ADPKD cells, blocked renal cyst formation in PKD mouse models, and reduced cyst formation and growth in a neonatal PKD mouse model ^26^. Although not identified in our analysis, STAT6, another member of STAT family, exhibits increased activity in two murine ADPKD models, mediated by interleukins IL4 and IL13 ^27^. Deletion of STAT6 in bpk/bpk, a model for ADPKD, resulted in less severe kidney disease and an improvement of kidney function. Pharmacological inhibition of STAT6 led to suppression of renal cystic growth, lower kidney weight and cystic index^27^. Given our observations and those previously reported for STAT6, we hypothesize that other STAT family members could also be involved in CHF/ARPDK pathogenesis.

IPA analysis predicted inhibition of SMAD7, a negative regulator of transforming growth factor β (TGF-β) signaling, in PCK rats at PND 30 and PND 90. Inhibition of SMAD7 suggests an increased activation of the TGF-β signaling pathway, which may further contribute to myofibroblast activation and fibrosis. Indeed, the anti-fibrotic role of SMAD7 was illustrated in multiple liver fibrosis models including the CCl^4^-induced liver fibrosis ^28^ and bile duct ligation models, as well as in isolated primary hepatic stellate cells, ^29^ and also in the unilateral ureteral obstructed (UUO)-induced renal fibrosis model ^30^. All these studies suggest that inhibiting TGF-β signaling through activating Smad7 can be a potential therapeutic tool in ARPKD.

In addition to the upstream regulators that were already established in ADPKD, such as the STATs and SMAD-TGFβ signaling, we also found some upstream regulators unique in ARPKD. MTPN (myotrophin) is a novel target predicted by IPA in ARPKD. Upstream analysis revealed that MTPN was activated from PND 15 to PND 90, suggesting its role is critical in ARPKD pathogenesis. Interestingly, MTPN stimulates insulin exocytosis, and its function can be suppressed by microRNA-375 (miR-375) ^31^, which is critical in maintaining normal glucose homeostasis, regulating insulin secretion, and pancreatic α- and β-cell turnover in response to increasing insulin demand in insulin resistance ^31, 32^. Interestingly, enhanced glycolysis, defective glucose metabolism, insulin resistance, and hyperinsulinemia have been reported in ADPKD patients ^33, 34^. Thus, our studies have identified MTPN as a novel therapeutic target in CHF/ARPKD, which may correct the metabolic disorder inherent in this disease.

Finally, our studies revealed inhibition of HNF4α, the master regulator of hepatic differentiation in CHF/ARPKD. HNF4α plays a critical role in liver development and hepatocyte differentiation. In fact, HNF4α is absolutely essential in differentiation of hepatocytes from hepatoblasts and its downregulation results in increased cell proliferation and cancer in the liver ^35, 36^. A detailed analysis of HNF4α target genes showed decline in several cytochrome P450 family members (*Cyp3a5, Cyp2c9, and Cyp2b6*), transporters (*Abcc2, Abcg5, Abcg8, Aqp8, and Slco1a1*), and lipid metabolism (*Acox2, Acsl1, Lipa, Pla2g4a, and Srebf1*). These data, for the first time reveal that CHF/ARPDK is not only associated with increased cystogenesis, fibrosis, and inflammation but also with significant dysregulation in hepatocyte differentiation and subsequent metabolic function, which may further increase disease-associated morbidity. Further studies are needed to investigate the role of HNF4α in CHF/ARPKD and how it regulates CHF/ARPKD pathogenesis.

Certain limitations of this study should be noted. First, we used whole liver for bulk RNAseq analysis. The observed gene expression changes, while novel and useful, include gene expression in all liver cell types (i.e. hepatocytes, cholangiocytes, liver sinusoidal endothelial cells, myofibroblasts, and any resident or infiltrating hepatic leukocytes). Future studies using single cell RNAseq are needed to determine cell-type specific gene expression changes in SD and PCK rats and in ARPKD patients compared to controls. Furthermore, a detailed proteomic analysis would be extremely useful. Whereas we confirmed upregulation of the top four genes increased throughout the time course, this candidate gene approach has its limitations that can be overcome by bulk or single cell proteomics.

In summary, this is the first comprehensive global hepatic gene expression analysis in a rat model of CHF/ARPKD over a prolonged pathogenic period. Our analysis provides strong evidence to support the ‘pathogenic triumvirate’ of proliferation-inflammation-fibrosis as underlying mechanisms driving cystogenesis and pericystic fibrosis in CHF/ARPKD. Further, our studies have revealed a critical therapeutic window between PND20 and PND30 prior to rapid disease progression which represents a previously unrecognized therapeutic opportunity. Future studies should determine if a similar therapeutic window exists in human CHF/ARPKD children. Moreover, we identified several novel therapeutic targets which could prevent disease progression within this therapeutic window and attenuate or halt progression of this devastating disease. Thus, these studies provide a solid foundation for launching further mechanistic investigations and drug development efforts to pharmacologically manage CHF/ARPKD in humans.

## Acknowledgements

These studies were supported by NIH: P20 RR021940, P20 GM103549, R01 DK98414. The Kansas Intellectual and Developmental Disabilities Research Center U54 HD 090216. The authors would like to thank Clark Bloomer and Roseann Skinner of the KUMC Genomics Core for the RNA sequencing. Dr. Satyajeet Khare is SPK is a beneficiary of a DST SERB SRG grant (SRG/2020/001414)

