## Supplementary material for "Global Transcriptomics of Congenital Hepatic Fibrosis in Autosomal Recessive Polycystic Kidney Disease using PCK rats"

Supplementary figure 1

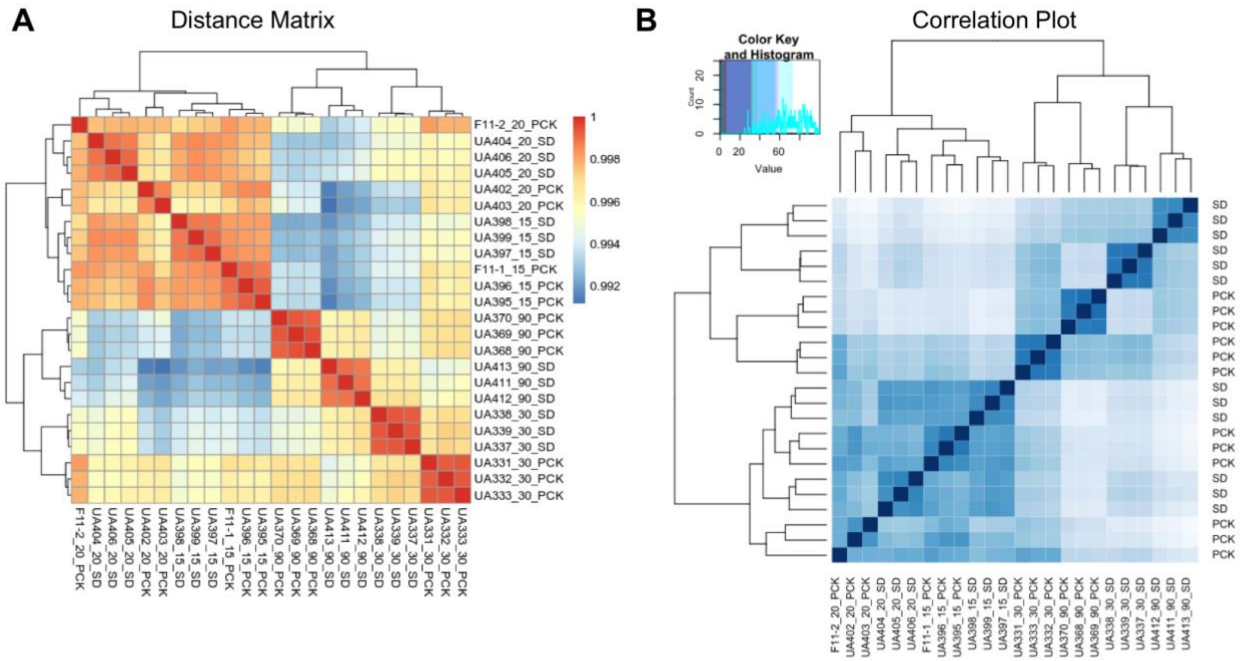

**Supplementary Figure 1. Data quality analysis of RNAseq data.** (A) Distance matrix plot and (B) Correlation plot show that biological replicates in each group cluster closer together indicating good samples quality, RNAseq analysis and overall good data quality.
